# Phase tracking algorithms detect both real and imaginary components of outer hair cell nonlinear membrane capacitance that exhibits dielectric loss

**DOI:** 10.1101/2022.01.01.474702

**Authors:** Joseph Santos-Sacchi

**Author notes:** ***Send correspondence to:*** Joseph Santos-Sacchi, Surgery (Otolaryngology), Neuroscience, and Cellular and Molecular Physiology, Yale University School of Medicine, BML 224, 333 Cedar Street, New Haven, CT 06510.

## Abstract

Outer hair cell (OHC) nonlinear membrane capacitance (NLC) represents voltage-dependent sensor charge movements within prestin (SLC26a5) that drive OHC electromotility. Dielectric loss, a shift in charge movement phase from purely “capacitive” to “resistive”, is likely indicative of prestin’s interaction with the viscous lipid bilayer and has been suggested to correspond to prestin power output. The frequency response of NLC in OHC membrane patches has been measured with phase tracking and complex capacitance methodologies. While the latter approach can directly determine the presence of dielectric loss by assessing charge movement both in and out of phase with driving voltage, the former has been suggested to fail in this regard. Here we show that standard phase tracking in the presence of dielectric loss does indeed register this loss. Such estimates of NLC correspond to the absolute magnitude of complex NLC, indicating that total charge movement regardless of phase is assessed, thereby validating past and present measures of NLC frequency response that limits its effectiveness at high frequencies. This observation has important implications for understanding prestin’s role in cochlear amplification.

## Introduction

Outer hair cell (OHC) membrane capacitance (C_m_) is composed of linear and voltage-dependent, nonlinear (NLC) parts (Ashmore, 1990; Santos-Sacchi, 1990, 1991). Both phase tracking methodology (Gale and Ashmore, 1997) and complex admittance assessment (Santos-Sacchi and Tan, 2020; Santos-Sacchi et al., 2021) have been used to estimate OHC NLC frequency response in membrane patches under voltage clamp. OHC NLC is a complex quantity (cNLC), separable into real and imaginary components that represent SLC26a5 (prestin) voltage-sensor charge movements differing in phase by 90 degrees with respect to driving membrane voltage (Santos-Sacchi and Tan, 2020; Santos-Sacchi et al., 2021). This separation results from dielectric loss, a frequency-dependent accumulation of charge that moves through a “resistive” path, typically generating heat. The imaginary component of NLC can inform on this behavior across frequency. Dielectric loss may have its basis in the interaction of prestin with the viscous plasma membrane where it resides. Recent cryo-EM studies, including our own, on prestin have found evidence of significant interactions of membrane lipid around and within protein helices (Bavi et al., 2021; Butan et al., 2021; Ge et al., 2021) that confirms expectations from earlier physiological studies that identified effects of membrane lipid modification (Santos-Sacchi and Wu, 2004; Sfondouris et al., 2008; Fang et al., 2010; Zhai et al., 2020).

Recently, Rabbitt (Rabbitt, 2020) claimed that all past measures of OHC NLC, and indeed all phase tracking C_m_ estimates in general, report only on the real component of complex capacitance. This is an important issue to clarify since it suggests that past measures of NLC have misjudged the OHC’s influence on cochlear amplification by neglecting an important component (dielectric loss) of prestin charge movement that may correspond to power output of electromotility. Indeed, it was questioned whether identified low-pass prestin kinetics is a limiting factor for OHC performance *in vivo* (Gale and Ashmore, 1997; Santos-Sacchi and Tan, 2018; Santos-Sacchi, 2019; Santos-Sacchi et al., 2019; Santos-Sacchi and Tan, 2019, 2020).

Here, we report that, in the presence of dielectric loss, measures of NLC employing phase tracking methods, provide the absolute 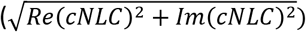, not real, value of complex capacitance. This indicates that total prestin charge movement, both in and out of phase with voltage, has been appropriately assessed in past studies. We demonstrate this with a standard electrical model with dielectric loss, with a 2-state kinetic model, and with macro-patch data from the OHC lateral membrane.

## Methods and Results

The most thorough method to measure membrane capacitance whether or not dielectric loss is present is that outlined by Fernandez et al. (Fernandez et al., 1982). In the absence of dielectric loss, only a real “capacitive” component of complex capacitance is present, whereas in its presence, both real and imaginary “resistive” components are present. Complex capacitance (cC_m_) is derived from the membrane admittance, *Y*_*m*_(*ω*), following removal of the influence of R_s_ and R_m_ from the input admittance, *Y*_*in*_(*ω*) of the simple model in **Fig 1**. Thus, complex capacitance is defined as

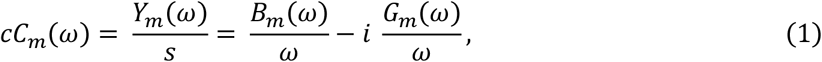

where *s=iω, ω* = 2*πf* and 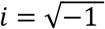.

**Fig. 1.**
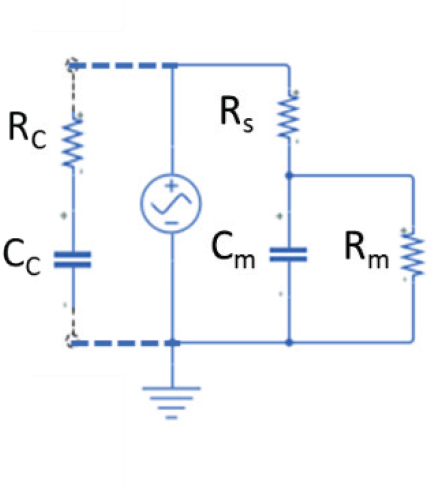
Phase detection implementation under voltage clamp. Capacitance compensation (R_C_ and C_C_) circuitry is added to the standard electrical model and the proper admittance phase angle and is determined to estimate membrane capacitance, C_m_. R_s_, electrode series resistance; R_m_, membrane resistance.

The complex components are a consequence of capacitive charge moving both in and out of phase with AC voltage, and Im(cC_m_) does *not* represent an actual membrane conductance.

A common method to measure membrane capacitance is the phase detection/tracking method (Neher and Marty, 1982; Fidler and Fernandez, 1989) which tracks changes in membrane current that correspond to changes in membrane capacitance. It is not clear how these measures relate to those obtained by complex capacitance estimation. Below we investigate the relationship.

### Phase detection/tracking method without dielectric loss

Interpretation of phase tracking membrane capacitance (C_m_) estimates is model dependent (**Fig. 1**). In the classical phase tracking method, we require to determine the proper phase angle to measure current that corresponds to changes in C_m_ (Neher and Marty, 1982; Fidler and Fernandez, 1989), i.e., the angle (β_ω_) of the partial derivative δY/δC_m_. The change in current is then factored by a gain (H_C_) of 1/abs(δY/δC_m_) to determine the actual capacitance change (see (Gillis, 1995)). β_ω_ can be determined exactly from the admittance at any two frequencies for the simple model of **Fig. 1** (Santos-Sacchi, 2004). However, for the single sine lock-in approach, the angle is estimated empirically. For example, following full capacitance compensation, where the admittance of the compensation circuitry (R_C_ and C_C_) exactly matches and is subtracted from the model admittance, a calibrated change in C_C_ can be used to estimate δY/δC_m_ and its angle. Alternatively, the angle of δY/δRs + pi/2 approximates that of δY/δC_m_ and can be empirically determined by dithering in a small change (ΔR_s_) in series resistance (**Fig. 1**). This angle can also be calculated from the input admittance of the model circuit, assuming the recording system is properly calibrated to account for extraneous phase and magnitude deviations (Zierler, 1992). At β_ω_, the real component of dither current is unresponsive to small changes in C_m_ and the imaginary output is then proportional to changes in C_m_, which is scaled by H_C_.

Analogously, we can work on the admittance of the model circuit. The input admittance at any frequency in the absence of capacitance compensation is

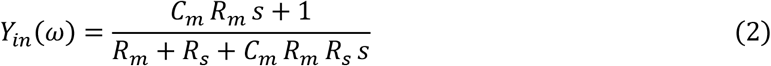

where *s=iω, ω* = 2*πf* and 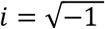.

Nulling Y_in_ with the two-component compensation circuitry requires that both R_C_ and C_C_ be adjusted to attain an input admittance (Y_C_) equal to that of the three-component model, Y_in_.

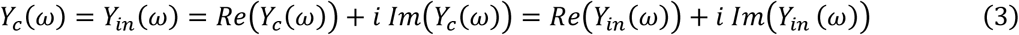

An equivalent increment in membrane capacitance, ΔC_m_, is obtained by slightly unbalancing C_C_.

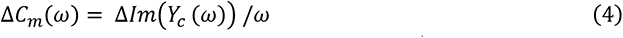

The resultant difference admittance, Y_ΔCm_ - Y_in_, provides an estimate of δY/δC_m_, its angle, β_ω_ and the gain factor, H_C_. Instead, H_C_ can be estimated from Y_in_ – see eq. 43 in (Gillis, 1995), and more exactly, β_ω_ can be determined from real (A_0_, A_1_) and imaginary (B_0_, B_1_) components of Y_in_ at any two frequencies (f_0_, f_1_) (Santos-Sacchi, 2004).

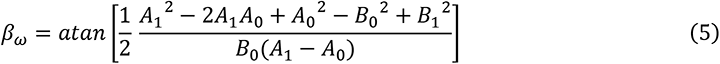

Using any of these approaches to determine proper inspection angle and gain provides essentially the same outcome that follows.

*Y*_*in*_(*ω*) is then monitored at its angle shifted by β_ω_ to give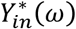, whose imaginary component factored by H_C_ provides any subsequent changes in C_m_.

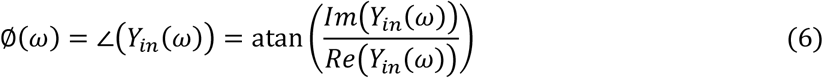

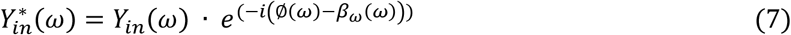

with the imaginary component corresponding to changes in C_m_.

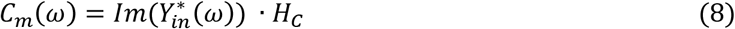

*C*_*m*_ estimated in this way is independent of stimulus frequency, as expected.

### Phase tracking with Dielectric loss

Dielectric loss is most simply modelled by inserting a resister (R_DiL_) in series with a capacitance (C_DiL_) (Fernandez et al., 1982). *R*_*DiL*_, *the equivalent series resistance (ESR), is hidden from physical inspection and cannot be removed, as it is intrinsic to C*_*DiL*_. **Fig. 2** illustrates the circuit used to model the addition of such loss to linear membrane capacitance, C_m_. The resistance results in a dissipation of power into the resistor, generating heat, and can be monitored with the imaginary component of complex capacitance, Im(cC_m_). What does standard phase tracking measure in the presence of dielectric loss? To mimic the addition of voltage-dependent OHC NLC, β_ω_ and H_C_ were first determined in the absence of dielectric loss. This would be equivalent to recording from OHCs at very positive membrane potentials (e.g., +160 mV) where NLC is lacking. Membrane capacitance is then tracked before and after the addition of a lossy capacitor in parallel with C_m_. When followed, the lock-in approach detailed above shows that the imaginary component of input admittance tracked at β_ω_, corresponds to Abs(cC_m_), thus providing an estimate of C_m_ across frequency that includes capacitor charge movement both in and out of phase with AC voltage. A Matlab script in the s-domain (**Fig. 3A, B**) confirms that phase tracking at most frequencies closely follows Abs(cC_m_). Note that at low frequencies Re(cC_m_) tracks Abs(cC_m_), but at high frequencies Im(cC_m_) tracks Abs(cC_m_). The cut-off frequency of Abs(cC_m_) depends inversely on the magnitude of R_DiL_ or dielectric loss.

**Fig. 2.**
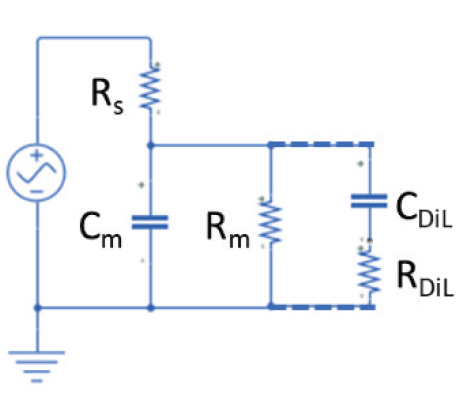
Dielectric loss is modeled by adding a lossy capacitor (C_DiL_ in series with R_DiL_) parallel to the membrane capacitance, C_m_.

**Fig. 3.**
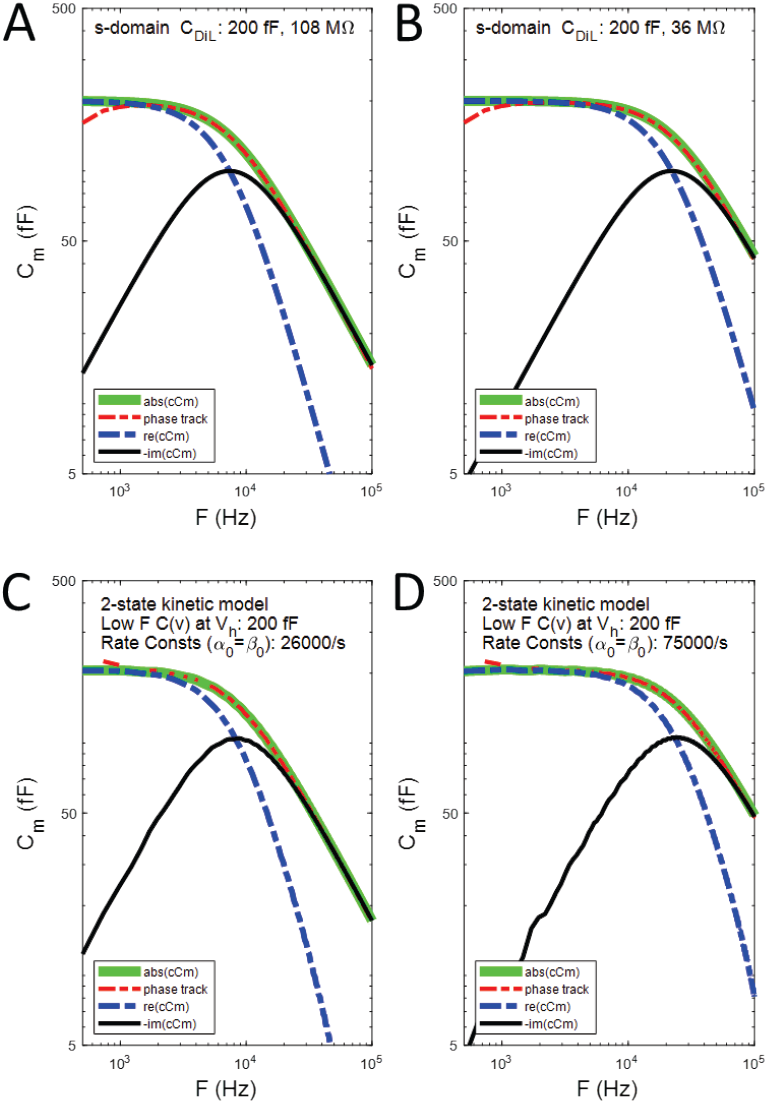
C_m_ estimates returned by complex and phase-tracking methods as a function of dielectric loss analyzed in the s-domain in Matlab (**A, B**) or as a function of 2-state kinetic transition rates analyzed in the time domain in Matlab Simulink (**C, D**) across frequency. Phase tracking provides estimates of Abs(cC_m_) across frequency and depends on the degree of dielectric loss or transition rates, thus including influences of charge movement in and out of phase with voltage. For the kinetic model, forward and backward rate constants are the same and linear capacitance is subtracted prior to plotting.

### Phase tracking with a 2-state voltage-dependent kinetic model

Fernandez et al. (Fernandez et al., 1982) pointed out that Eyring rate models exhibit dielectric loss characteristics. Consequently, real and imaginary components of voltage-dependent capacitance are generated in a simple 2-state kinetic model of prestin (Santos-Sacchi et al., 2021), where expanded (X) and compact (C) state populations redistribute during changes in membrane voltage.

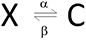

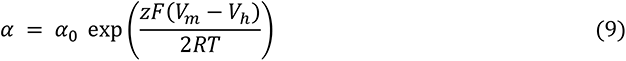

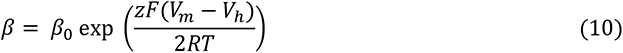

Both forward and backward rates (α, β, respectively) are governed solely by membrane voltage (V_m_) about a characteristic potential, V_h_, where charge is distributed equally on either side of the membrane field. *z* denotes the unitary particle charge (e^-^ X distance travelled perpendicular to the membrane field). F, R and T have their usual meanings. V_h_ is set to zero mV.

As above, measures are made relative to a holding voltage where voltage-dependent capacitance is absent, i.e., +200 mV. Here, again (**Fig. 3C, D**), phase tracking estimates at the voltage (V_h_) of peak C_m_(v) correspond to measures of Abs(cC_m_(v)) across frequency, with cut-off frequencies of Abs(cC_m_(v)) increasing with the particular transition rate constants (26000/s or 75000/s), with α_0_ = β_0_. The frequency cut-off at V_h_ is given by 1/ (2π τ), with τ=1/ (α_0_ + β_0_).

### Voltage-dependent nonlinear capacitance in OHC membrane patches

Voltage-dependent nonlinear capacitance (NLC) arises from restricted sensor charge movement (displacement currents) within membrane bound, voltage-dependent proteins (Fernandez et al., 1982; Santos-Sacchi, 1991; Kilic and Lindau, 2001). We find that in OHCs, complex NLC (cNLC) presents significant dielectric loss, i.e., an Im(cNLC) component, in measures of prestin (SLC26a5) generated capacitance (Santos-Sacchi et al., 2021). We model/fit NLC as a molecular capacitor obeying Boltzmann statistics (Santos-Sacchi, 1991).

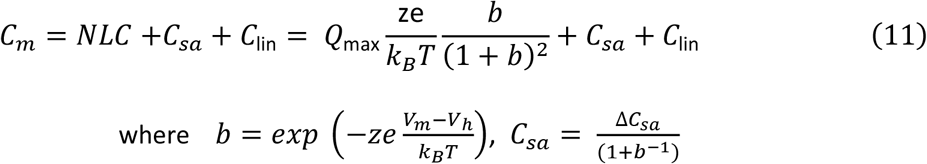

Q_max_ is the maximum nonlinear charge moved, V_h_ is voltage at peak capacitance or equivalently, at half-maximum charge transfer, V_m_ is membrane potential, *z* is valence, C_lin_ is linear membrane capacitance, e is electron charge, *k*_*B*_ is Boltzmann’s constant, and T is absolute temperature. C_sa_ is a component of capacitance that characterizes sigmoidal changes in specific membrane capacitance, with ΔC_sa_ referring to the maximal change at very negative voltages (Santos-Sacchi and Navarrete, 2002; Santos-Sacchi and Song, 2014).

**Fig. 4A-D** shows components of complex NLC measured in lateral membrane macro-patches of the OHC as a function of holding voltage and frequency. Data sets are combined and reanalyzed from our recent publications (Santos-Sacchi and Tan, 2019, 2020; Santos-Sacchi et al., 2021). In **Fig. 4E**, fitted NLC components measured near V_h_ (from **Fig. 4C, D**) are plotted across frequency, as well as NLC_Vh_ estimates determined from the phase tracking approach detailed above (*eq. 3-8*), relative to measures at +160 mV holding potential. Phase tracking estimates were smoothed by running average across 35 indices at 24 Hz resolution for frequencies above 10 kHz. As with model evaluations, it can be seen that phase tracking estimates of NLC (green line) correspond to Abs(cNLC) (blue downward triangles).

**Fig. 4.**
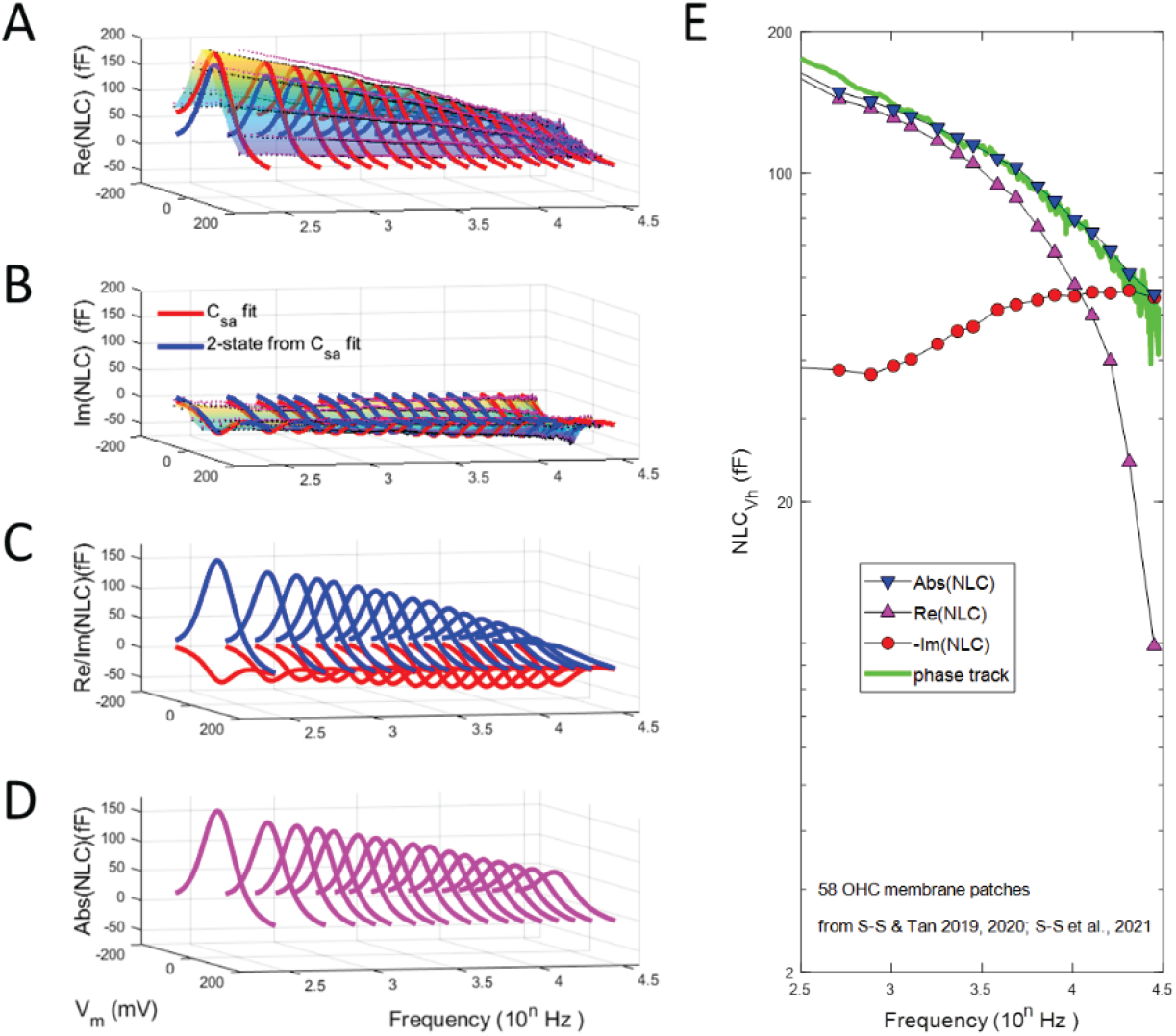
Real and imaginary components of macro-patch NLC with comparisons of Abs(cNLC) and phase tracking estimates. **A)** Real component of complex NLC across frequency. Blue dots are mean, red dots are +SE. Fits are made to *eq*.*11* to extract the 2-state (blue line) response at selected frequencies. **B)** Imaginary component of complex NLC. Magnitude increases with frequency, with less than a full reciprocal trade-off with real NLC. **C)** Plot of the 2-state fits. **D)** Plot of the absolute magnitude of NLC, i.e., 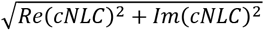, showing a continuous reduction of prestin activity across frequency. **E)** Plots of NLC near V_h_ (−40 mV). Real NLC is much lower pass than Abs(cNLC). Notably, phase tracking estimates of NLC (green line) correspond to those of Abs(cNLC). Phase tracking results were obtained from admittance analysis of averaged raw currents. Complex NLC was determined as previously described (Santos-Sacchi et al., 2021).

## Discussion

Phase tracking techniques that have been used to measure OHC NLC in patches evaluate to close estimates of Abs(cNLC), representing total charge moved by prestin’s voltage sensors, both in and out of phase with voltage. Indeed, the frequency response of Re(cNLC) and Abs(cNLC) differ, with Abs(cNLC) being wider in bandwidth. Note that at low frequencies Re(cNLC) tracks Abs(cNLC), but at high frequencies Im(cNLC) is predicted by modelling to track Abs(cNLC), indicating that Abs(cNLC) measures fully characterize the low-pass behavior of Im(cNLC) at very high frequencies. Thus, past frequency response measures of OHC NLC have not missed contributions from dielectric loss. In fact, the phase tracking data of Gale and Ashmore (Gale and Ashmore, 1997) and the Abs(cNLC) data of Santos-Sacchi and Tan (Santos-Sacchi and Tan, 2020) are in good agreement, each consistent with reductions in OHC NLC as a power function of frequency. It should also be noted that measures of Abs(cNLC), which phase tracking provides, cannot be used to predict the underlying real and imaginary components since we have found that the frequency response of real and imaginary NLC components are individually susceptible to membrane tension, despite stability of Abs(cNLC) across frequency (Santos-Sacchi et al., 2021).

Why does phase tracking return Abs(cNLC)? The technique strives to minimize the real component of admittance (or current) by rotating its measurement angle, thus shifting any real component into the imaginary component detector. Rotating the angle of any complex number in order to zero the real component will give Im(A+B i) = Abs(A+B i). The detection of the Abs(cNLC) value is thus an unintended quirk of the method.

To summarize, we have shown in simple models and in biophysical data that NLC assessment with phase tracking methods provide close estimates of Abs(cNLC), i.e., 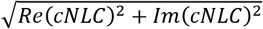, not Re(cNLC), as previously claimed (Rabbitt, 2020). Consequently, prestin performance has been realistically appraised during the last few decades, and its low-pass characteristic remains a limiting factor for the protein’s influence on cochlear amplification at very high acoustic frequencies. Finally, should dielectric loss be present in any cell’s membrane capacitance, for example as expected from channel gating charge movements, estimates of vesicle release by C_m_ phase tracking may be impacted in a manner not previously appreciated; that is, C_m_ estimates will be influenced not only by capacitive gating charge components (Horrigan and Bookman, 1994) but also by resistive gating charge components.

## Acknowledgments

This research was supported by NIH-NIDCD R01 DC016318 and R01 DC008130. We thank Fred Sigworth for comments on the manuscript.

